# On the Analysis of Protein Genetic Architecture: Response to “Protein sequence landscapes are not so simple”

**DOI:** 10.1101/2024.09.17.613512

**Authors:** Yeonwoo Park, Brian P.H. Metzger, Joseph W. Thornton

## Abstract

We recently reanalyzed 20 combinatorial mutagenesis datasets using a novel reference-free analysis (RFA) method and showed that high-order epistasis contributes negligibly to protein sequence-function relationships in every case. Dupic, Phillips, and Desai (DPD) commented on a preprint of our work. In our published paper, we addressed all the major issues they raised, but we respond directly to them here. 1) DPD’s claim that RFA is equivalent to estimating reference-based analysis (RBA) models by regression neglects fundamental differences in how the two formalisms dissect the causal relationship between sequence and function. It also misinterprets the observation that using regression to estimate any truncated model of genetic architecture will always yield the same predicted phenotypes and variance partition; the resulting estimates correspond to those of the RFA formalism but are inaccurate representations of the true RBA model. 2) DPD’s claim that high-order epistasis is widespread and significant while somehow explaining little phenotypic variance is an artifact of two strong biases in the use of regression to estimate RBA models: this procedure underestimates the phenotypic variance explained by RBA epistatic terms while at the same time inflating the magnitude of individual terms. 3) DPD erroneously claim that RFA is “exactly equivalent” to Fourier analysis (FA) and background-averaged analysis (BA). This error arises because DPD used an incorrect mathematical definition of RFA and were misled by a simple numerical relationship among the models that only holds only for the simplest kinds of datasets. 4) DPD argue that using a nonlinear transformation to account for global nonlinearities in sequence-function relationships is often unnecessary and may artifactually absorb specific epistatic interactions. We show that nonspecific epistasis caused by a limited dynamic range affects datasets of all types, even when the phenotype is represented on a free-energy scale. Moreover, using a nonlinear transformation in a joint fitting procedure does not underestimate specific epistasis under realistic conditions, even if the data are not affected by nonspecific epistasis. The conclusions of our work therefore hold: the genetic architecture of all 20 protein datasets we analyzed can be efficiently and accurately described in an RFA framework by first-order amino acid effects and pairwise interactions with a simple model of global nonlinearity. We are grateful for DPD’s commentary, which helped us improve our paper.

## Introduction

In a recently published paper, we reanalyzed 20 combinatorial mutagenesis datasets using a novel reference-free analysis (RFA) method and showed that protein sequence-function relationships are surprisingly simple: first-order effects and pairwise interactions between amino acids, along with a simple global nonlinearity, account for virtually all of phenotypic variance in all datasets (Park et al. 2024). Dupic, Phillips, and Desai (DPD) commented on the first preprint of our paper, arguing that RFA is “exactly equivalent” to existing formalisms and that higher-order interactions, despite contributing little to phenotypic variance, are “widespread and significant” (Dupic *et al*. 2024). They also argued that our modeling of global nonlinearity artifactually simplified some datasets, and ultimately that protein sequence-function relationships are “not so simple.”

We appreciate DPD’s engagement with our work. The published version of our paper addresses all of their arguments, but here we directly respond to each point, referring to relevant parts of our paper and providing further analyses.

### 1. RFA is distinct from reference-based analysis (RBA) and is more robust to model misspecification and measurement noise

DPD argue that RFA is “exactly equivalent” to RBA coupled with regression. Their primary rationale is that when RFA and RBA models of the same order are estimated from the same data by regression, the two models predict the same phenotype for any genotype. This equivalence arises because regression optimizes the model parameters to minimize the prediction error regardless of whether the true values of the parameters actually minimize the error. All genetic models of the same order therefore converge on the same set of predicted phenotypes under regression. If the sole purpose of analysis were to predict the phenotype of unseen genotypes, all genetic models would indeed be equivalent in DPD’s sense.

We have much more in mind, however. The purpose of using a genetic model to interpret DMS data is to reveal the causal relationships between a protein’s sequence and its biochemical phenotype, a fundamental form of knowledge in genetics and biochemistry. The genetic model quantifies the phenotypic effects of individual amino acids and their interactions. These model terms then enable macroscopic descriptions, like the fraction of phenotypic variance explained by subsets of the model’s effects, such as those at each epistatic order or at particular sequence site(s) of interest. The genetic model can be interpreted in the context of protein structure, providing biophysical insights into how the sequence determines the protein’s phenotype. And the model can provide evolutionary understanding, by revealing how particular elements of the genetic architecture cause the distribution of phenotypes across connected sequence space and thereby affect evolutionary trajectories (Weinreich et al. 2006; Palmer *et al*., 2015; Jalal *et al*. 2020; Metzger *et al*. 2024). Different genetic models decompose the sequence-function relationship in mathematically different ways and therefore lead to different inferences about genetic architecture, its causes, and its evolutionary implications. The terms of the RFA model represent the global effects of amino acid states across all genotypes, whereas those in RBA represent the effects of mutations on a particular reference sequence. These differences also have practical consequences for inferring the model from experimental data. When a truncated RBA model is estimated by regression, the inferred values of individual terms and the variance explained at each order can be wildly incorrect even when the data are perfect, because of an inherent misfit between the structure of the RBA model and the optimization criterion regression uses to estimate the model; these estimation problems are exacerbated by measurement noise. Not recognizing this artifact led DPD to the contradictory claim that high-order interactions can be “widespread and significant” while contributing negligibly to phenotypic variance. RFA does not suffer from this problem and can accurately characterize the genetic architecture from noisy and incomplete data. Below we explain these points with explanation and examples.

#### RFA and RBA conceptualize the sequence-function relationship differently

The terms of the RBA and RFA models represent distinct causal factors, differ in mathematical definition, and map differently to phenotype. These fundamental differences do not depend on the method used to estimate the terms of the models or the variance that subsets of those terms explain. In this subsection, we summarize the different parametrizations; for more detail, see Fig. 1 and 2 and Supplementary Section 1.1 in our paper.

**Fig. 1.**
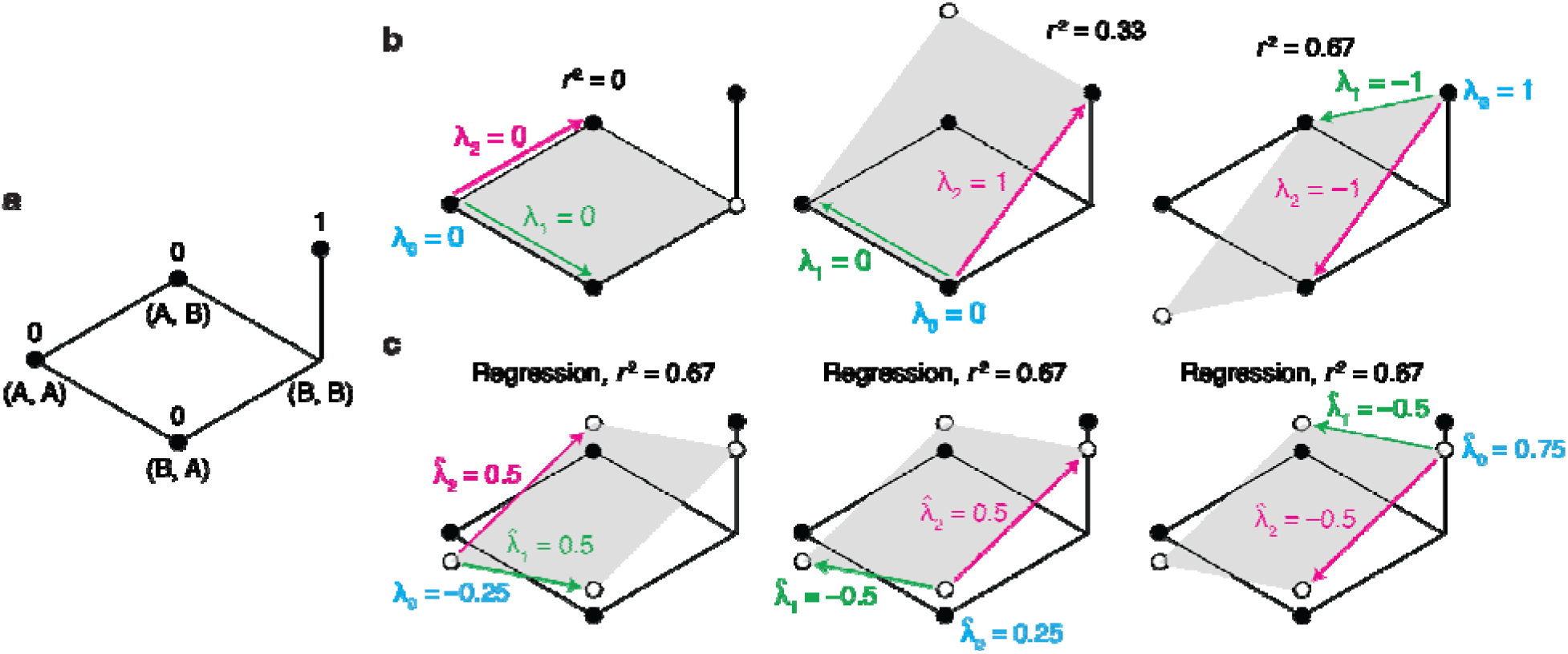
Pathologies of RBA-regression. **a**, Genetic architecture consisting of two states (A and B) in two sites. The four possible genotypes are arranged on a plane, with their phenotype indicated by the height of the dot. **b**, True first-order RBA models, using genotype (A, A), (B, A), or (B, B) as wild-type. For each case, the zero-order term (λ_0_) and the two first-order terms (λ_1_ and λ_2_) are shown. The grey plane is the genetic architecture predicted by the first-order model, with the fraction of phenotypic variance explained (*r*^2^) indicated. **c**, Fitting the first-order RBA model by regression. The inferred effects and the fraction of phenotypic variance explained are shown. These figures have been excerpted from our paper.

**Fig. 2.**
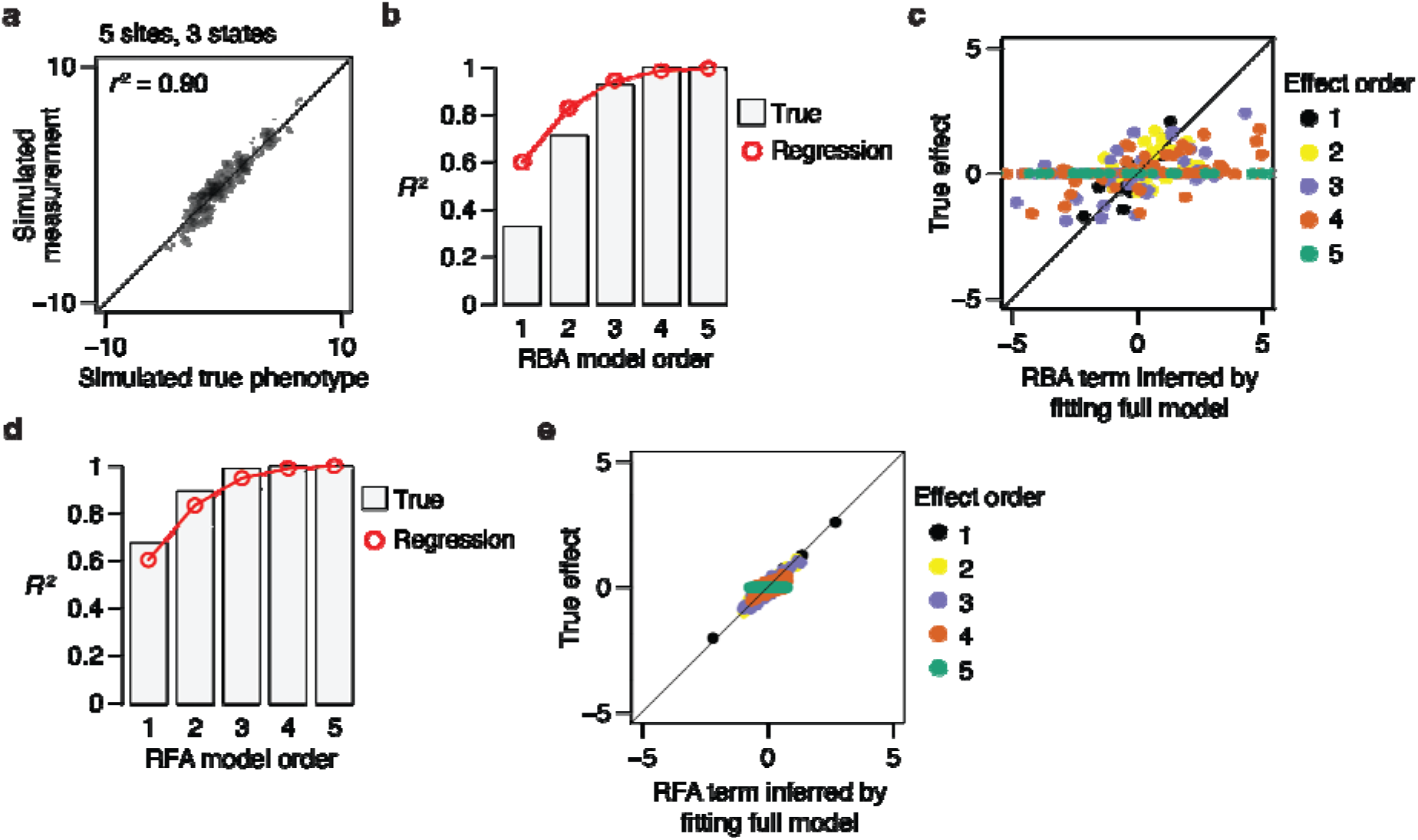
RBA-regression mischaracterizes the genetic architecture. **a**, Genetic architecture with 5 sites and 3 possible states was simulated by drawing reference-based effects from the standard normal distribution (but setting all fifth-order effects to zero). A small amount of simulated noise was added to the resulting phenotypes. **b**, Fraction of phenotypic variance explained by the true and regression-estimated RBA models. **c**, RBA terms estimated by fitting the fifth-order model, compared with their true values. **d**, Fraction of phenotype variance explained by the true and regression-estimated RFA models for the same simulated data. **e**, RFA terms estimated by fitting the fifth-order model, compared with their true values.

The purpose of the RFA model is to explain a protein’s global genetic architecture – to quantitatively describe how amino acids and their interactions determine the phenotype across the entirety of sequence space. Each first-order RFA term represents the average effect of an amino acid at a sequence position, defined as the difference between the mean phenotype of the subset of genotypes sharing that amino acid and that of all genotypes. Each second-order term represents the mean interaction of two states at a pair of sites, defined as the difference between the mean phenotype of the subset of genotypes sharing the two states and the expected phenotype given the two first-order effects. Higher-order terms follow this pattern, representing the mean deviation of variants sharing a specific set of states from the expectation based on lower-order terms.

The purpose of RBA is to provide a locally anchored view of genetic architecture by quantifying the effects of mutations and their interactions when introduced into a designated wild-type sequence. Each first-order RBA term represents the effect of mutating the wild-type amino acid at one site into a different state; it equals the difference between the phenotype of the one protein that contains just that mutation and the wild-type. Each second-order term is the difference between the phenotype of a double-mutant and that expected given the two single-mutants. Higher-order terms continue this pattern, representing the deviation of one higher-order mutant from the expectation given the lower-order mutants on the path to the wild-type.

These structural differences between the models lead to very different representations of the sequence-function relationship and different inferences about genetic architecture. For example, RFA quantifies the average causal role of amino acid *states*, whereas RBA quantifies the role of *mutations* from a wild-type state to another when introduced into a particular genetic background. RFA models the effect of all states contained in the library, but in RBA, the wild-type states have no explicit effect on the phenotype, because – not being mutations -- they do not appear in the model at all. RFA effects are averaged over all genetic backgrounds, but RBA terms are fundamentally conditioned on the wild-type background: a first-order RBA term for a mutation incorporates not only the effect of the mutant state but also the interactions of that state with the wild-type state at every other site in the protein, as well as the loss of the wild-type state and all its interactions. Because RBA expresses each epistatic effect as the deviation of an individual mutant genotype relative to individual lower-level mutant genotypes, any idiosyncracy in the neighborhood of the wild-type sequence will propagate into epistatic effects with increasing distance from that reference. As a result, RBA almost always attributes a greater causal role to epistasis, especially at high orders, than RFA does. And reassigning the wild-type sequence necessarily alters the values of RBA terms and the fraction of phenotypic variance explained by each epistatic order; therefore, the apparent complexity of the genetic architecture varies depending on the choice of wild-type sequence. By contrast, there is a single RFA description for each genetic architecture, and this description minimizes the phenotypic variance attributed to epistasis across all of sequence space.

RFA and RBA therefore represent fundamentally different ways of conceptualizing genetic architecture and explaining how any particular protein’s sequence determines its phenotype. These differences pertain no matter what method is used to estimate the terms of each model.

#### Coupling RBA with regression causes biased and anomalous inferences

There are two ways in which RBA models have been estimated from experimental data. One is to directly compute each term exactly according to its definition as the difference between the phenotypes of particular mutant genotypes. The other, which is the focus of DPD’s argument, is to estimate all the terms jointly from all the phenotypes using regression. Several publications, mostly from Desai’s group, have followed this approach, which proceeds in two steps. First, the fraction of phenotypic variance explained by each epistatic order is estimated by fitting a series of truncated RBA models to the entire data. Second, the values of individual terms are estimated by fitting the complete untruncated model (or a truncated model statistically as powerful as the complete model).

DPD claim that RFA and RBA are “exactly equivalent” when estimated by regression; specifically, they claim that the two approaches are “exactly as robust to measurement noise and missing data and lead to identical results.” In fact, the two models behave very differently and yield distinct inferences when estimated by regression.

RFA is designed so that the individual terms and the fraction of variance explained by each epistatic order can be accurately estimated by fitting truncated models using regression. Specifically, least-squares regression computes the parameter values that minimize the sum of squared error (SSE) between the model-predicted and measured phenotypes, with the sum taken across all genotypes in the dataset. In an RFA model truncated at any order, the true RFA terms also minimize the SSE across sequence space, because they are defined relative to the mean phenotypes in subsets of that space. The regression-based estimates of RFA terms are therefore accurate, converging on the true values as long as sampling is sufficient and unbiased. RFA estimates are also robust to using truncated models: each order of RFA terms produces an orthogonal pattern of phenotypic variation—with variation caused by higher-order terms appearing as noise around the phenotypes predicted by lower-order terms—so the inference of truncated models is not biased by the exclusion of higher-order terms. Our paper shows that the regression-estimates of RFA terms are unbiased and robust to measurement noise, converging rapidly on the true values even when sampling density is low (Figs. 1 and 3 and Supplementary Section 1.1).

**Fig. 3.**
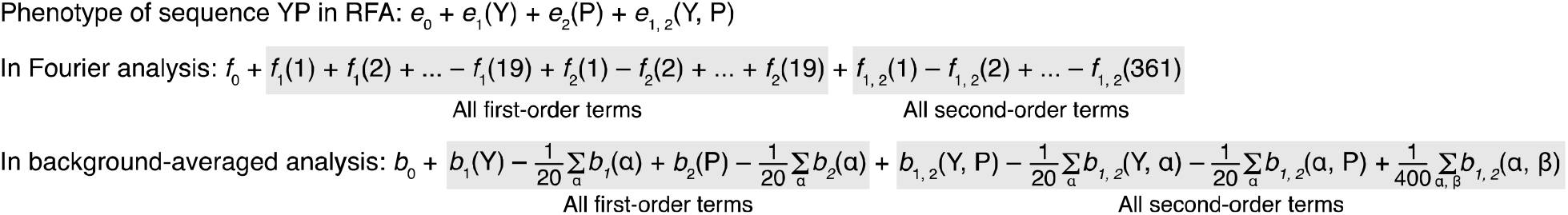
Mapping of RFA, FA, and BA coefficients to phenotype. The mapping is shown for a two-amino-acid sequence YP. Subscript 0 indicates the zero-order term; subscript 1 or 2, first-order term for site 1 or 2; subscript 1, 2 indicates the pairwise epistatic term.

By contrast, both stages of RBA-regression cause strong biases. The first part of RBA-regression overestimates the fraction of phenotypic variance explained by the true RBA model. This bias has already been established in the literature (Otwinowski and Plotkin, 2014), and exemplifies a general bias that arises whenever regression is used to fit uncentered interaction models where variables are correlated across orders. RBA-regression is biased because RBA terms are defined in ways that conflict with the optimization criterion of regression. In the true RBA model, low-order terms exactly describe the corresponding low-order mutants and do not attempt to describe any higher-order mutant. But regression minimizes the SSE across all sampled genotypes. Therefore, when a truncated RBA model is fit by regression, the low-order terms are forced to fit the phenotypic variation caused by the higher-order terms excluded from the model, resulting in a biased model that explains more phenotypic variance than the true RBA model explains at that order. The predicted phenotypes match those predicted by RFA, because these are the phenotypes that minimize SSE, and in turn they explain the same fraction of variance as the RFA model truncated at the same order. These results accurately represent the true RFA genetic architecture but are a biased representation of the true RBA architecture.

Consider the simple genetic architecture in Fig. 1a below. The true zero-order RBA term is defined as the wild-type phenotype, but regression using a truncated zero-order model estimates it to be the mean phenotype of all genotypes, which offers a lower SSE than does the wild-type phenotype and in turn explains more variance than the true zero-order RBA model does. Similarly, the true first-order terms exactly fit the difference between each single mutant and the wild-type, but regression of a truncated first order model finds values that minimizes the SSE across the entire data set, thus explaining more phenotypic variance than does the true model does (compare Fig. 1b and c). The estimates of individual terms are also typically inflated in magnitude (Figs. 1b, c). In a larger genetic architecture with more sites, this bias would continue with each order, systematically overestimating the variance explained by each truncated model until the complete model is reached. A related pathology of RBA-regression is that it infers the same variance partition no matter what wild-type sequence is chosen, whereas the true RBA variance partition depends on the choice of reference sequence (Fig. 1c).

In the second stage of RBA-regression, a different bias arises. Here the values of individual terms are estimated by fitting the complete model to the data. Because the model is complete, regression is gratuitous, and the regression-estimated terms are identical to directly computing each term according to its definition. When measurement noise is present in the data, the RBA epistatic terms estimated by regression are therefore wildly inflated, especially at high orders, as demonstrated in our paper (Fig. 2) and already established in the literature (Poelwijk et al. 2016).

Fig. 2 below illustrates the pathological behavior of RBA-regression caused by these two biases. We generated an RBA genetic architecture with no epistatic effects above the fourth order and then simulated phenotypic data using this architecture with a small amount of measurement noise (Fig. 2a). Regression systematically overestimates the variance explained by each truncated RBA model, with the first-order model accounting for more than double its true *R*^2^ (Fig. 2b). When the complete model is fit, there appears to be pervasive high-order epistasis, including fifth-order effects that dwarf all lower-order effects, even though the true value of all fifth-order terms is zero (Fig. 2c). These twin biases together lead to the anomalous impression that high-order epistasis terms are widespread and large but somehow explain virtually no phenotypic variance. By contrast, the RFA variance partition and model terms are estimated accurately (Fig. 2d, e).

These results contradict DPD’s claim that RBA-regression is “exactly as accurate and robust to measurement noise and partial sampling as the reanalysis of Park et al.” RBA-regression terms and variance partition are biased towards overestimating and underestimating the complexity of the genetic architecture under the RBA model, respectively. RFA-regression is unbiased and converges on accurate estimates under the RFA model even when sampling is sparse and measurements are noisy.

#### Misinterpretation of phenotype predictions, variance partition, and transformability of model terms

RBA-regression and RFA yield the same predicted phenotypes and variance partition across orders, but this fact is a property of regression itself; it provides no evidence that the underlying models or the inferences drawn from the inferred models are equivalent. RBA-regression misrepresents the phenotypes and variance partition associated with the model that is putatively being fit. The results obtained happen to correspond to the true values associated with a different model – RFA. Regression-based results are meaningful and accurate only if the RFA model is the one being estimated; they are misleading if they are associated with the RBA model.

DPD claim that the “equivalence” of RBA-regression and RFA “does not only hold at the level of model performance or of predicted phenotypes: all of the coefficients inferred in either framework are also exactly interchangeable.” They base this statement on the fact that the terms of RBA and RFA models are related by a set of linear transformations. This point is technically correct, but its significance is overstated. Both RBA and RFA are linear models, each of which represents all phenotypes in a dataset as a sum over the relevant effect coefficients in the model. Because the degrees of freedom are the same, the complete set of terms in an RFA model can therefore be transformed into the terms of the RBA model through an elaborate set of weighted addition and subtraction operations, and vice versa. Essentially, this operation involves taking all the terms of one model, predicting all the phenotypes, and then decomposing those phenotypes again into the terms of the other model. For example, obtaining just the zero-order RFA term from the RBA model requires a transformation involving every single term at every order of the entire RBA model. The fact that a complicated transformation is required to remap one mathematical decomposition of the sequence-function relationship onto the other does not provide evidence that the models are equivalent. Rather, it highlights their fundamental differences and explains why they respond so differently to regression, truncation, and propagation of measurement error.

#### DPD’s claim that protein sequence-function relationships are “not so simple” arises from the biases of RBA-regression

DPD do not dispute our finding that high-order epistasis contributes negligibly to phenotypic variance in all 20 datasets, but they contend that this observation does not imply that protein sequence-function relationships are simple, because “high-order epistasis can be widespread and significant while still not substantially increasing variance explained.” This paradoxical statement arises from their reliance on RBA-regression. Fitting truncated RBA models by regression underestimates the variance explained by high-order RBA terms, but fitting the full model dramatically inflates the magnitude of individual high-order terms. The result is the illusion that high-order terms are widespread and of large magnitude while somehow contributing negligibly to phenotypic variance.

These considerations rebut DPD’s claim that our development of RFA was gratuitous and that using RBA-regression would have reached “the exact same conclusion.” We found that protein sequence-function relationships are shaped almost entirely by first- and second-order determinants as defined by the RFA formalism. The terms at these orders are consistently of substantial magnitude while higher-order terms are generally very small (see Supplementary Fig. 5 of our paper). If RBA-regression were used instead, the conclusions would be very different and also incorrect: that phenotypic variation when analyzed from the RBA perspective is attributable mostly to low-order determinants (which is false in many datasets), that the variance partition does not depend on the wild-type sequence chosen (which is almost always false), and that high-order RBA terms are large but somehow explain little variance (which is self-contradictory).

#### The utility of RBA and RFA

We stress that although the RBA formalism is not suited to estimating a protein’s global genetic architecture, it has a legitimate use: to understand the effects of mutations in the local neighborhood of a particular sequence of interest. In this case, low-order terms can be directly computed, although high-order terms must be interpreted with care because even small measurement noise can snowball with increasing distance from the wild-type. RBA models should never be estimated by regression, however, because of the anomalous and inaccurate results that this approach yields.

DPD state that interactions between mutations and their implications for evolutionary trajectories are captured more directly by RBA coefficients than by RFA. The terms of the RBA model are indeed structured to express the effects of mutations in the reference sequence, but RFA can provide a deeper analysis of the effects of mutations on particular backgrounds, and it can do so for any neighborhood in sequence space, not only the wild-type. In the RFA model, introducing a mutation into a particular background exchanges the existing amino acid state for a new state, changing the first-order effects at this site and all of the epistatic interactions that this site has with every other site in the protein at every order. The terms of the RFA model allow the phenotypic effects of the mutation to be decomposed into all these contributing factors. In this way, RFA can reveal in detail how the underlying genetic architecture determines the effects of particular mutations in particular backgrounds and how this architecture affects evolutionary trajectories across any part of sequence space (see, for example, Metzger et al. 2024).

Finally, DPD claim that RBA models are more useful for analyzing protein phenotypes because RFA terms lack biochemical meaning. In fact, the terms of the two formalisms have equal relevance to biophysical quantities; it is only the genetic parameterization that differs. Consider a mutational dataset that accurately measures the free energy change (ΔG) of binding to some ligand. The zero-order term of RBA is the ΔG of binding by the reference protein; the zero-order term in RFA is the average ΔG of binding by all genotypes. A first-order RBA term is the difference in ΔG between the protein that carries an amino acid mutation and the wild-type protein; the first-order RFA term is the average difference in ΔG between genotypes carrying that state and all genotypes. The RFA terms are thus no less or more biochemically meaningful than the RBA terms.

### 2. RFA is distinct from “Hadamard methods”

DPD argue that RFA is also “exactly equivalent” to “Hadamard-based models.” By “Hadamard-based models,” DPD refer to both Fourier analysis (FA) and background-averaged analysis (BA). All three methods are in fact conceptually and mathematically distinct, and the differences matter because they affect the methods’ interpretability and robustness. Our paper compares the methods in detail (Fig. 3 and Supplementary Sections 1.2 and 1.3). Here we summarize the essential differences and suggest potential sources for DPD’s error.

#### Differences among the formalisms

In formal terms, the methods differ in two critical ways: how genotypes are decomposed into model coefficients, and how the coefficients combine to predict the phenotype of each genotype (Fig. 3 below). In RFA, each term is the average effect of an amino acid state or a combination. Using *q* to denote the number of possible states, there are *q* first-order terms at each site, *q*^2^ second-order terms at each pair of sites, and *q*^*k*^ terms for each *k*-tuple of sites. The phenotype of any genotype is a simple sum of the terms that correspond to the amino acid states and combinations in the genotype; terms for any other states make no contribution to the predicted phenotype.

FA is structured differently. FA terms do not directly represent the effects of amino acids. Rather, each amino acid is recoded as a series of (1, –1) coordinates in a Fourier space with (*q –* 1) dimensions, implemented by using a specific kind of Hadamard matrix or graph Fourier bases. A first-order term in the FA model represents the average effect of having a 1 or –1 in one of the Fourier dimensions relative to the global average, not the effect of any amino acid. Similarly, the (*q –* 1)^2^ second-order terms represent the pairwise interactions among Fourier terms, not amino acids. Unlike RFA, the phenotype of any genotype is a sum over every single term in the entire model, with unique signs applied to each term depending on the particular genotype (Fig. 3 below).

Like FA, BA also employs (*q* − 1)^*k*^ terms at each order *k*, but they are defined differently from FA. A first-order BA term represents the effect of mutating a reference state to a mutant state at one site, averaged across all sequence backgrounds at other sites. Similarly, a second-order BA term represents the interaction of two mutations at a pair of sites averaged across all sequence backgrounds at other sites. Although the BA terms themselves have an intuitive meaning, the phenotype of any genotype is a signed and weighted sum of all the terms in the entire model, including those that represent mutations not in the genotype of interest (Fig. 3 above). This complicated structure arises because the terms represent phenotypic differences from the reference state, but the model’s intercept is the global average, not a reference sequence as in RBA. The encoding of genotypes into BA terms and the signs and weights are again derived from a Hadamard matrix, but the matrix is different from that used for FA. Examples of each matrix and their differences are presented in Supplementary Sections 1.2 and 1.3 of our paper.

These differences among the models have practical consequences. First, they affect interpretability: RFA terms have a straightforward genetic meaning, directly expressing the average phenotypic effects of amino acids and their interactions, and the simple mapping of terms to the phenotype directly reveals how any genotype’s phenotype arises from its sequence states. Second, the methods perform differently in the face of partially sampled and noisy data. Each RFA term is averaged over a larger number of measurements than each BA term, so the error on estimated terms is smaller for RFA. Because the FA and BA phenotype is a sum over every term in the model, the error associated with each term propagates to all genotypes. As a result, RFA results in more accurate estimation and/or prediction than BA and FA, particularly when the number of states is large (see Fig. 3 of our paper).

#### DPD’s incorrect claim of equivalence among the three formalisms appears to have two sources

First, DPD’s formal definition of RFA is incorrect. They define RFA in their Eq. (1), which expresses the phenotype of any genotype as a uniquely signed sum over every term in the model, with the terms being multiplied by either 1 or –1 depending on the particular genotype. This is not RFA. In the correct definition of RFA, each phenotype is a simple sum over only the terms for the amino acid states and combinations present in the genotype of interest; that is, RFA uses (1, 0) rather than (1, –1) encoding. Rather than showing RFA, DPD’s Eq. (1) actually corresponds to FA in the special case of *q* = 2; DPD’s derivations on p. 5 confirm that Eq. (1) is indeed FA. Incorrectly defining RFA using the definition for FA may have led DPD to the erroneous belief that RFA and FA are identical.

A second source of error is that DPD consider only binary state space (*q* = 2), but many mutational datasets examine many possible states, up to all 20 amino acids. DPD argue incorrectly that “RFA and Hadamard methods are identical… parameterized by exactly the same coefficients (appropriately re-indexed and scaled).” In the special case of *q* = 2, the terms of the models—although they are different in mathematical definition and conceptual meaning—do have a simple numerical relationship, which allows them to be easily interconverted with a change of sign and scaling. For example, let *e*(A) and *e*(B) be the RFA first-order terms for the two states at a site, *f* be the first-order Fourier term (encoding state A as –1 and B as 1), and *b* be the background-averaged mutational effect from state A to B. The following equality holds by virtue of the terms’ definition: –*e*(A) = *e*(B) = *f* = *b*/2.

But this simple relationship does not hold when *q* > 2. In this case, the terms of the models can be interconverted only through a very complicated set of equations (see Supplementary Sections 1.2 and 1.3 in our paper). When there are 20 states, for example, each first-order RFA term is a signed sum across all 19 first-order FA terms, each second-order RFA term is a signed sum across all 361 second-order FA terms, each third-order RFA term is a signed sum across all 6,859 third-order FA terms, and so on. The interconversion between BA and RFA (and between BA and FA) are similarly complex, but in that case the signs are different and each term in the equation is weighted by its order. The fact that the terms of these formalisms can be interconverted does not mean they are equivalent; rather, it underscores how differently they define and represent the sequence-function relationship.

### 3. Benefits of modeling nonspecific epistasis outweigh potential costs

A key finding in our paper is that the bounding of measurable phenotype by upper or lower limit creates strong nonspecific epistasis in most of the datasets we analyzed; incorporating this bounding into the sequence-function model using a sigmoid link function allows low-order genetic models to explain virtually all of phenotypic variance. DPD question the benefits of this procedure and raise concern about potential costs. We agree that these are important issues to consider, but we explain below why we disagree with DPD’s assessment on both counts.

#### Benefits

Nonspecific epistasis (NSE) arises when the effects of all mutations on the phenotype can be mapped to the same nonlinear relationship. For example, consider a case in which mutations act independently but combine on a multiplicative scale, but the phenotype is analyzed in absolute rather than relative terms. The absolute effect of a mutation that increases the phenotype by 10% in every background will nevertheless depend on the phenotype of the variant in which it is measured. If the genetic architecture is modeled without accounting for this nonlinearity, pervasive specific interactions among amino acids must be invoked to explain the context-dependence of mutations, yielding a gratuitously complex description of the genetic architecture. If the nonlinearity can be incorporated into the analysis, however, the underlying simplicity of mutations’ specific effects can be accurately revealed. This is a well-established issue in the literature (e.g., Otwinowski et al. 2018).

##### Benefits of the sigmoid link function

In our paper, we assessed a particular approach to incorporating nonspecific epistasis: using a sigmoid link function in a joint fitting procedure with the RFA model of specific interactions. Although many different link functions could be used, we explored the sigmoid function because it is mathematically simple and experimentally motivated. There is always a limited dynamic range over which variation in a phenotype can be accurately measured, and in many datasets a substantial fraction of variants have phenotypes outside this range. The bounding of measurable phenotype may arise for biological reasons—such as saturation of the biological mechanisms that produce the phenotype—or for technical reasons—because experimental assays have lower and upper bounds of detection, near which further phenotypic reductions or increases, respectively, can no longer be observed. We know of no assays that lack such bounds.

In our paper, we showed that incorporating nonspecific epistasis using the sigmoid link function has large and general benefits. Specifically, we established that: 1) Using this model substantially improves the fit to 17 of our 20 experimental datasets compared with excluding the link function; 2) Incorporating nonspecific epistasis simplifies the inferred specific genetic architecture in all of these cases, often to a large extent; 3) The improvement in fit is tightly correlated with the fraction of genotypes that have phenotypes outside the dynamic range; and 4) Simulations show that when phenotypes are generated with phenotype bounds and with specific epistasis, the link function effectively captures this phenomenon and allows the RFA variance partition to be accurately inferred. We conclude that whenever bounds affect the phenotype—as they do in most datasets—using the sigmoid link function improves the inference of genetic architecture.

DPD argue that when “a substantial fraction of sequences lead to nonfunctional proteins (with phenotypes below the experimental threshold of detection), … this is often indicative of strong high-order epistasis rather than a limited dynamic range” The logic of this statement is flawed. If many sequences are below the threshold of detection, this fact alone establishes that the dynamic range is limited and many sampled genotypes are outside that range. Those conditions are sufficient to cause nonspecific epistasis, because the phenotypic effect of deleterious states in these backgrounds is masked by the nonlinearity imposed by the lower bound. DPD assert instead that the clustering of many proteins with phenotypes at the lower bound indicates pervasive specific epistasis, but this conclusion does not necessarily follow: this same pattern can instead be produced by a limited dynamic range and a simple genetic architecture in which many amino acid states impair function, thus causing the majority of sequences to have low phenotypic values. This kind of architecture is likely to be quite common, particularly in multistate datasets; a recent publication provides an empirical example (Metzger et al. 2024). We observed that in most datasets, many genotypes are clustered near minimum and/or maximum phenotypic values; this provides prima facie evidence of nonspecific epistasis but not of high-order specific epistasis.

##### Free energy phenotypes

DPD question whether nonspecific epistasis always has benefits. They assert that “when the measured phenotypes are a thermodynamic state variable (e.g., free energy of binding), it is natural to expect that they behave additively without any such transformation.” In practice, this expectation does not hold. If free energy effects were known exactly, then using them as the modeled phenotype would indeed remove one important source of global nonlinearity—the Boltzmann distribution, which expresses how effects on free energy of a biophysical state determine the fractional occupancy of that state and, in turn, any phenotype that is proportional to occupancy.

We are unaware, however, of any combinatorial mutagenesis dataset that directly measured free energy, nor do we know of any assays that can measure free energy or its proxies without detection limits. Instead, free energy effects are generally estimated indirectly using an assay that measures a molecular phenotype, such as fluorescence, that indicates occupancy of some state (e.g., fraction of proteins ligand-bound), across a range of conditions (e.g., ligand concentration); free-energy effects are estimated by fitting biophysical models to these data. Assays of this type always have a limited dynamic range, below which the phenotype cannot be distinguished from background signal and measurement noise, and above which changes in phenotype are no longer apparent. Near the bounds, measured phenotypic differences between variants underestimate their actual differences, leading to nonlinearity in observed occupancies and hence nonlinearity of the inferred free energies. Unless all variants in the library are within the linear dynamic range, nonspecific epistasis will affect the free energies.

In the empirical data we analyzed, expressing phenotypes in terms of free energy does not eliminate nonspecific epistasis caused by phenotype bounding (Fig. 4 above). There are 9 datasets in which the phenotype is expressed as free energy of binding; in 7 of these, many variants lie outside the dynamic range of measurement, and in each case the genetic architecture can be explained more simply when NSE is incorporated using the sigmoid link function. In fact, 2 of the 3 datasets with the largest fraction of variants outside the dynamic range express the phenotype on a free energy scale.

**Fig. 4.**
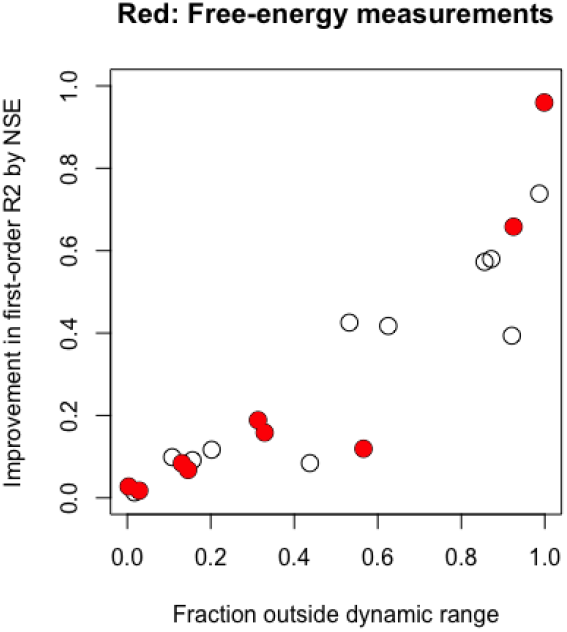
The sigmoid link function captures the NSE arising from limited dynamic range of measurement. Improvement in *R*^2^ of the first-order RFA model conferred by the sigmoid link is shown for each dataset (dot) against the fraction of genotypes at the lower or upper bound. Red, datasets in which the phenotype is expressed on the free energy scale.

##### The case of CR9114-H1

DPD argue that the CR9114-H1 antibody-antigen pair is an example in which there is no nonlinearity. To the contrary, the details of this case underscore the value of incorporating phenotype bounds using the sigmoid link function. In this experiment, binding of variants of the antibody CR9114 to influenza H1 was assessed using a yeast-surface display assay with a fluorescently tagged ligand across a range of concentrations; the fluorescence measurements were then fit to a binding curve to estimate the dissociation constant (K_D_), and the genetic architecture was modeled using –logK_D_ as the phenotype. DPD claim that all the data are in the dynamic range of measurement and that no link function should be used, but the data do not support this interpretation.

We analyzed these data (Fig. 5 below), and inferred substantial nonspecific epistasis due to phenotype bounding, with an upper bound –logK_D_ of 9.5. The second-order RFA model including the sigmoid link function explains 95% of genetic variance, but excluding the link function reduces this to 87%, leaving almost three times more variance to be explained by higher-order epistasis. More than 30% of all phenotypes are clustered in a peak around – logK_D_ of 9.5, with many variants at or below this threshold and none substantially above it. This pattern is precisely as expected if this is the top of the dynamic range with some measurement noise around it. DPD argue that the dynamic range is not limited at 9.5, because when the same technique is applied to a different antibody-antigen pair (CR6261-H9), some variants have estimated –logK_D_ over 10. This argument is not valid, however. Different antibody-antigen pairs have different affinities, expression levels, thermodynamic stabilities, propensities to be degraded in the cell, propensities to be displayed on the cell surface, etc., any of which can cause differences in the upper-bound phenotype that can be produced and observed in the assay. It is highly implausible that, if the upper bound in the CR9114-H1 dataset were indeeds greater than 10, that so many genotypes would be clustered around 9.5 and not one of the ∼64,000 genotypes would have a phenotype at or above 10.

**Fig. 5.**
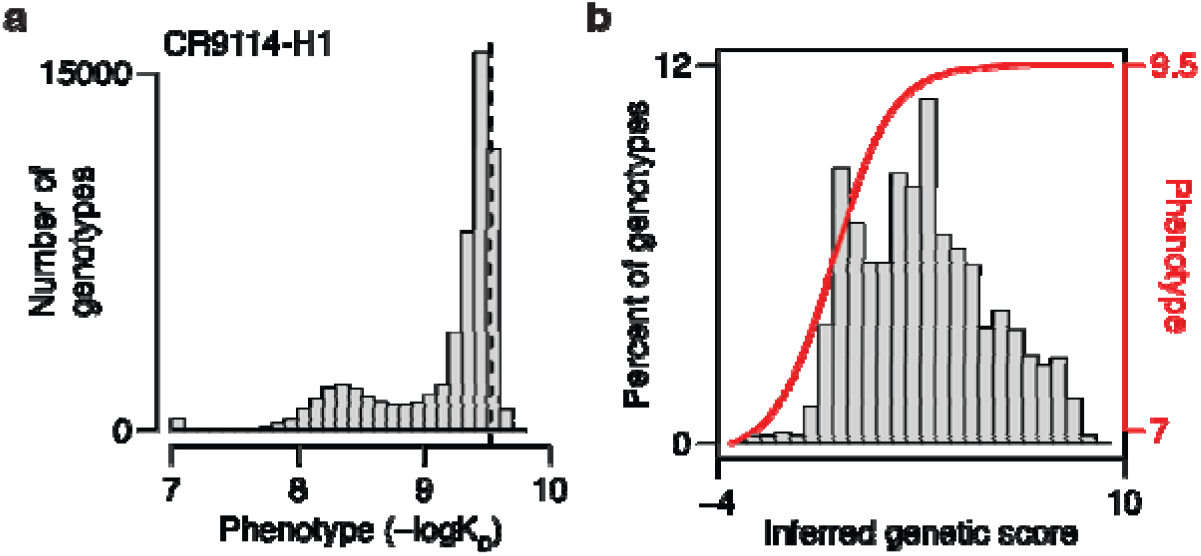
Upper bounding in the CR9114-H1 dataset. **a**, Distribution of phenotype. Dashed line marks the upper bound inferred by our sigmoid link (–logK_D_ = 9.5). DPD argue that the true upper bound is beyond 10, but none of the 2^16^ (= 65,536) genotypes have a phenotype at or beyond 10. **b**, Second-order RFA with the sigmoid link. Histogram shows the distribution of inferred genetic score across genotypes, and the red curve shows the inferred link function. 33% of genotypes have a genetic score that is masked by upper bound. The *R*^2^ of second-order RFA is 0.95, which drops to 0.87 when the sigmoid link is not used.

Given all these considerations, we believe that our original conclusion holds: incorporating NSE using a sigmoid transformation has substantial benefits when analyzing genetic architecture from most combinatorial mutagenesis experiments—those in which many variants are near the apparent limits of the dynamic range. There are a few datasets in which most variants are within the dynamic range and the benefits are small, but the best way to assess this possibility is to model the genetic architecture using a joint fitting procedure.

#### Costs

It is important to weigh these benefits against potential disadvantages of incorporating NSE. A legitimate concern is that using a nonlinear transformation might obscure specific epistasis, absorbing real amino acid interactions into a spurious model of nonspecific epistasis. Our approach is designed to minimize this problem by jointly fitting the sigmoid model of NSE and RFA models, rather than transforming the data first and then inferring specific effects. But whether this approach is effective—and understanding the potential impacts if it is not—merits consideration.

##### Absorbing specific epistasis into the nonlinear transformation

DPD argue that “specific (‘idiosyncratic’) epistatic effects can always be captured by a global transformation, and global epistasis can always be represented by specific epistatic coefficients. The question then becomes: which representation of the genotype-phenotype architecture is more meaningful?” Nonspecific epistasis can indeed always be misinterpreted as the result of a large number of specific genetic interactions, because a complete model of specific epistasis can be made to exactly fit any set of sequence-phenotype associations.

But the converse is not true, at least for the sigmoid link function. There are many forms of specific epistasis that cannot be absorbed by the sigmoid model, which is highly constrained in shape and the forms of nonlinearity it can therefore fit. The sigmoid transformation can accommodate only data patterns in which the effect of a mutation has a larger magnitude when introduced into genetic backgrounds within the dynamic range than it has near the phenotypic bounds. It can never account for sign epistasis, for example, or for nonmonotonic or bell-shaped relationships. For specific interactions to produce a pattern that can be entirely absorbed by the sigmoid transformation’s upper bound, amino acids with positive first-order effects must interact with each other strongly enough that their negative interactions exactly cancel out the first-order effects. To counterfeit a lower bound, amino acids with negative first-order effects must interact positively with each other. To counterfeit a sigmoid relationship with both upper and lower bounds—as seen in many of the datasets we analyzed—both of these patterns must occur in the same dataset. These scenarios seem very unlikely, particularly given our observation that epistatic terms are small relative to first-order effects in every dataset we examined—even those in which the sigmoid transformation has little or no effect on the model fit.

##### Empirical conditions: the case of CH65-MA90

The key question is the degree to which specific epistatic interactions are absorbed by using the sigmoid link function under realistic conditions. In our paper, we addressed this issue using an empirically derived simulation in which no nonspecific epistasis is present but the sigmoid link function is applied. In our analysis of the protein-antibody pair CH65-MA90, virtually all genotypes are within the dynamic range of measurement, suggesting very little nonspecific epistasis. We estimated the RFA model for this dataset with no sigmoid link function and then used the distribution of coefficients at orders 1, 2, and 3 to repeatedly simulate genetic architectures and associated phenotypes; a small amount of measurement noise was included but no nonspecific epistasis.

We fitted RFA models with and without the sigmoid link function to these data to see if the link function would absorb the specific epistasis (Supplementary Fig. 2 in our paper). We found that variance partition across orders is estimated accurately, and these inferences are unaffected by whether or not the link function is used. The minimum and maximum of the sigmoid function are estimated to be far beyond the range of phenotypic prediction, so the transformation has no effect. Even when the true generating model contains only third-order epistatic effects, all variance is correctly attributed to third-order epistasis (Fig. 3 of our paper). Specific epistasis is therefore not absorbed by the link function, even when nonspecific epistasis is absent.

##### DPD’s simulation

DPD assembled a simulation in which specific epistatic interactions produce a “noticeable nonlinearity” in a plot of the relationship between observed phenotypes and that predicted by a first-order model. This visual analogy says nothing about whether using a link function to incorporate NSE would in fact obscure these interactions. To address this question, we generated phenotypic data under DPD’s simulation conditions and analyzed them with and without the sigmoid link function. We note that these are unrealistic conditions—second-order interactions are huge (on average 6 times greater than first-order effects), they are all negative, there is no nonspecific epistasis, and the data are very noisy.

When we analyzed the phenotypes generated by this architecture, we found no difference between the fraction of variance explained by the second-order RFA model whether or not NSE is incorporated, and the inferred epistatic terms are also identical and are correctly estimated (Fig. 6 below). The truncated first-order model does yield a slight overestimate of the variance caused by first-order effects, but this problem disappears when second-order terms are included in the model. Within the fitted second-order model, the relative contribution of first- and second-order terms corresponds to the true variance partition. Moreover, if the ratio of the average magnitude of second-order to first-order effects is reduced to 2 instead of 6, the variance partition estimated using the truncated models becomes accurate.

**Fig. 6.**
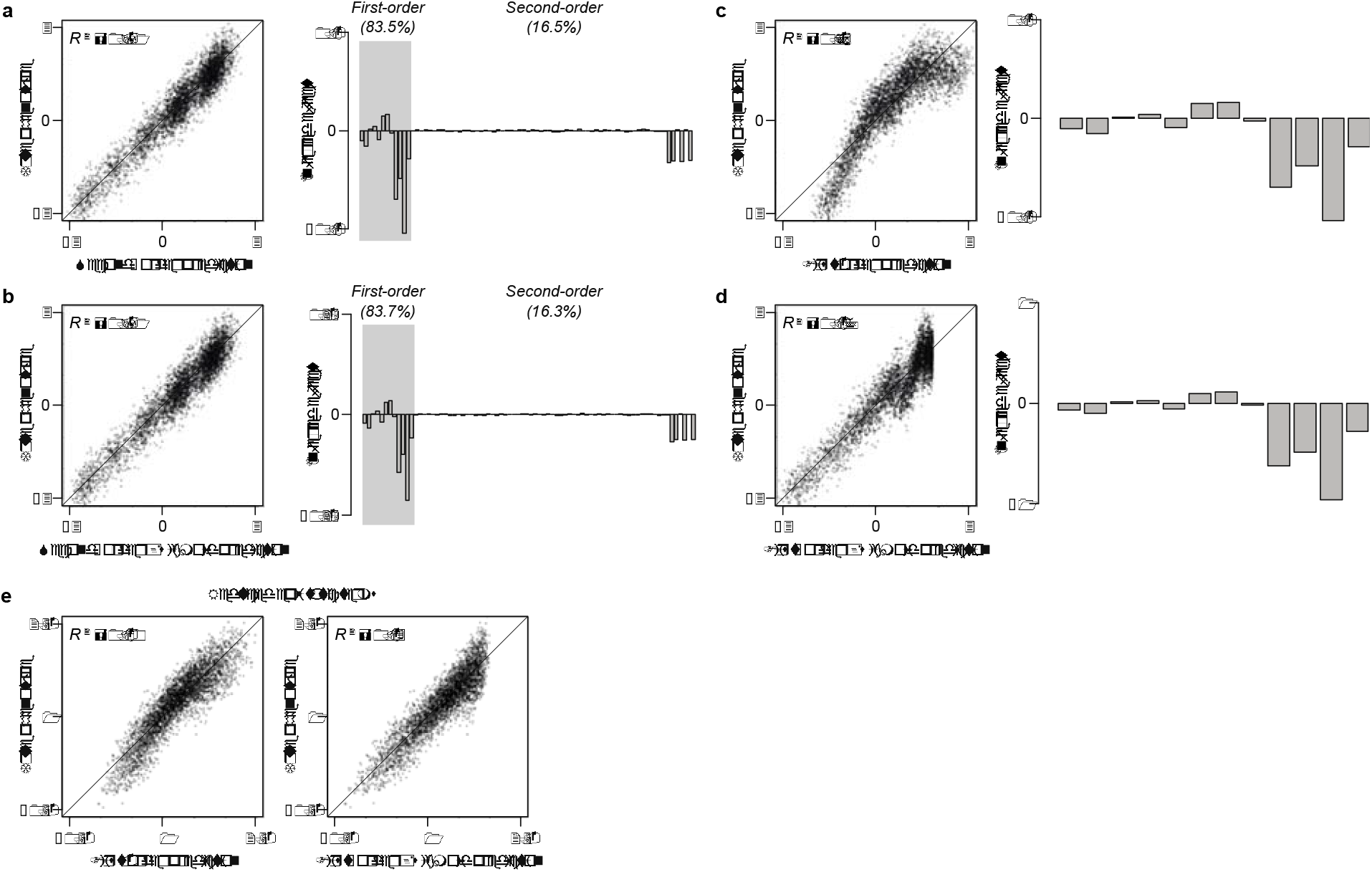
Re-evaluating DPD’s simulation of NSE. Following the conditions described in DPD, we simulated a 12-site, 2-state genetic architecture, sampling the intercept and first-order effects from *N*(2, 1) and *N*(0, 0.2^2^), respectively, and setting the pairwise interactions for site-pairs (9, 10), (9, 11), (10, 11), and (11, 12) to –1 and all others to zero. The fraction of variance explained by the first- and second-order RFA is 83 and 17%, respectively. Although DPD do not specify the amount of noise in the simulation, we noticed that their Fig. 3A involves substantial measurement noise, so we added noise to account for 10% of phenotypic variance. **a**, Second-order RFA without modeling NSE. The predicted vs. simulated phenotype (left) and the inferred RFA terms (right) are shown. Despite the noise, the relative variance contribution of first- and second-order terms within the inferred model (83.5 and 16.5%) corresponds to the true ratio. **b**, Second-order RFA with the sigmoid link. The predicted phenotypes, inferred terms, and the relative variance contribution of first- and second-order terms are virtually identical to those obtained without modeling NSE. (The scale of inferred terms is different because of the presence and absence of sigmoid link.) **c**, First-order RFA without modeling NSE. **d**, First-order RFA with the sigmoid link. The variance explained is slightly overestimated, although the inferred terms are virtually identical (save for the different scale). **e**, We modified DPD’s simulation by setting the pairwise interactions to –1/3 instead of –1. Shown are first-order RFA without or with the sigmoid link. The true fraction of variance attributable to first-order effects is 89% (80% after accounting for noise).

Taken together, these analyses indicate that under empirically derived conditions, the sigmoid link function does not underestimate the importance of specific epistasis. Under particular extreme and unrealistic conditions, variance partitioning using truncated RFA models and a link function has the potential to underestimate the complexity of genetic architecture. However, the estimated genetic architecture itself will be inferred accurately, as long as the important epistatic terms are present in the model, and the fitted model can be used to accurately estimate the variance partition.

#### Costs versus benefits

The benefits of incorporating nonspecific epistasis can therefore be weighed against the costs. When a dataset is affected by nonspecific epistasis, incorporating those effects into the model will result in the most compact, efficient, and accurate description of the genetic architecture. This will prevent inferring a gratuitously elaborate model of rampant specific epistatic interactions to explain why the effect of mutations differ among backgrounds. Because all assays have limited dynamic range and in most datasets the measured phenotypes are affected by these bounds, the benefits of incorporating NSE are expected to accrue widely or even universally. The extent of this benefit will depend on the fraction of variants that are near the limits of the dynamic range.

The costs, by contrast, appear to be small. When nonspecific epistasis is weak or absent, our data indicate that the joint fitting procedure is expected to yield accurate results as long as the relevant specific epistatic terms are included in the model. It remains possible that there could be particular scenarios in which unmodeled specific epistasis could partially counterfeit a sigmoid relationship and reduce a truncated model’s power to identify specific interactions. But there is no evidence in any of the empirical or simulated datasets that this occurs under realistic conditions. Even if it did, use of the complete RFA model allows epistatic terms and their variance contribution to be accurately assessed.

We conclude that the benefits of using the link function outweigh the costs. The benefits apply almost universally and are often very large in providing a more compact and efficient description of the genetic architecture of real proteins. The potential costs appear to be small and, based on present knowledge, are expected to occur very infrequently under realistic conditions. Further research in this area, however, is warranted.

## Conclusion

Based on the above considerations, the claims of our paper hold: the genetic architecture of all 20 protein datasets we analyzed can be efficiently and accurately described by first-order amino acid effects and pairwise interactions with a simple model of global nonlinearity in an RFA framework. We believe that RFA presents a useful new way to quantitatively dissect proteins’ genetic architecture, and that it can be effectively coupled with a simple representation of the effect of phenotypic bounding to yield a compact, efficient, and accurate description of proteins’ sequence-function relationships. As we discuss in the published paper, however, there is much further work to be done to evaluate the genetic architecture of different kinds of protein functions and to assess the methods available for this purpose.

DPD’s critique provided an impetus for us to improve our paper by addressing and clarifying these important issues. We are grateful for their engagement with our research and for all their work to advance the analysis of protein genetic architecture.

## Acknowledgements

This work was supported by National Institutes of Health grants R35-GM145336 (J.W.T), R01-GM131128 (J.W.T.), R01-GM121931 (J.W.T.), F32GM122251 (B.P.H.M.), and by a Samsung Scholarship (YP).

## References

Dupic, T., Phillips, A.M., and Desai, M.M. (2024). Protein sequence landscapes are not so simple: on reference-free versus reference-based inference. bioRxiv 2024.01.29.755800.

Jalal et al. (2020). Diversification of DNA-Binding Specificity by Permissive and Specificity-Switching Mutations in the ParB/Noc Protein Family. Cell Rep. 32: 107928.

Metzger, B.P.H., Park, Y., Starr, T.N., and Thornton, J.W. (2024). Epistasis facilitates functional evolution in an ancient transcription factor. eLife 12: RP88737.

Otwinowski, J. and Plotkin, J.B. (2014). Inferring fitness landscapes by regression produces biased estimates of epistasis. Proc. Natl. Acad. Sci. USA 111: E2301–E2309.

Otwinowski, J., McCandlish, D.M., and Plotkin, J.B. (2018). Inferring the shape of global epistasis. Proc. Natl. Acad. Sci. USA 115: E7550–E7558.

Palmer, A.C., Toprak, E., Baym, M., Kim, S., Veres, A., Bershtein, S. and Kishony, R. (2015). Delayed commitment to evolutionary fate in antibiotic resistance fitness landscapes. Nat. Commun. 6, 7385.

Park, Y., Metzger, B.P.H., and Thornton, J.W. (2024). The simplicity of protein sequence-function relationships. Nat. Commun 15, 7953 (2024).

Poelwijk F.J., Krishna, V., and Ranganathan, R. (2016). The Context-Dependence of Mutations: A Linkage of Formalisms. PLoS Comput. Biol. 12: e1004771.

Weinreich, D.M., Delaney, N.F., Depristo, M.A. and Hartl, D.L. (2006). Darwinian evolution can follow only very few mutational paths to fitter proteins. Science 312, 111–114.

